# Utility of the *Onchocerca volvulus* mitochondrial genome for delineation of parasite transmission zones

**DOI:** 10.1101/732446

**Authors:** Katie E Crawford, Shannon M Hedtke, Stephen R Doyle, Annette C Kuesel, Samuel Armoo, Mike Osei-Atweneboana, Warwick N Grant

## Abstract

In 2012, the reduction in *Onchocerca volvulus* infection prevalence through long-term mass ivermectin distribution in African meso- and hyperendemic areas motivated expanding control of onchocerciasis (river blindness) as a public health problem to elimination of parasite transmission. Given the large contiguous hypo-, meso- and hyperendemic areas with an estimated population of 204 million, sustainable elimination requires an understanding of the geographic, and in turn genetic, boundaries of different parasite populations to ensure interventions are only stopped where the risk of re-introduction of the parasite through vector or human migration from areas with ongoing transmission is acceptable. These boundaries, which define the transmission zones of the parasite, may be delineated by characterising the parasite genetic population structure within and between potential zones. We analysed whole mitochondrial genome sequences of 189 *O. volvulus* adults to determine the pattern of genetic similarity across three West African countries: Ghana, Mali, and Côte d’Ivoire. Population structure measures indicate that parasites from the Pru, Daka and Black Volta/Tombe river basins in central Ghana belong to one parasite population, showing that different river basins cannot be assumed to constitute independent transmission zones. This research forms the basis for developing tools for elimination programs to delineate transmission zones, to estimate the risk of parasite re-introduction via vector or human movement when mass ivermectin administration is stopped in one area while transmission is ongoing in others, to identify the origin of infections detected post-treatment cessation, and to investigate whether migration contributes to persisting prevalence levels during interventions.

## 1. Introduction

Onchocerciasis, or river blindness, is a disease caused by the parasitic filarial nematode, *Onchocerca volvulus*, and transmitted by *Simulium* spp. blackflies, whose larvae develop in fast-flowing streams and rivers. The high level of onchocerciasis-related morbidity (in particular blindness, visual impairment, and skin disease) and resulting social and economic burden on communities led to the implementation of large-scale control and elimination programs. An estimated 45 million people would have been infected world-wide in 2014 without these programs (Remme et al., 2017). The World Health Organization (WHO) estimated that in 2017, 205 million people globally should be included in control and elimination programs, with over 99% (204 million) in sub-Saharan Africa (World Health Organization, 2017).

Mass drug administration of ivermectin (MDAi) has been the principal strategy for control of onchocerciasis since 1987, after MDAi was demonstrated to be safe and Merck & Co, Inc. (Kenilworth, NJ, USA) decided to donate ivermectin (Mectizan^®^) (De Sole et al., 1989). The Onchocerciasis Elimination Programme for the Americas (OEPA) targeted elimination of morbidity and transmission through biannual MDAi from its initiation in 1993, and the African Programme for Onchocerciasis Control (APOC, 1995-2015) targeted control of onchocerciasis as a public health problem through annual MDAi in areas with an infection prevalence exceeding around 40%, a prevalence which was previously shown to be associated with an increased risk of blindness (Prost et al., 1979; UNDP/World Bank/WHO Special Programme for Research and Training in Tropical Diseases, 1992). Research studies and epidemiological evaluations (Diawara et al., 2009; Fobi et al., 2015; Traore et al., 2012) suggested that annual or biannual MDAi may have interrupted parasite transmission in many areas in Africa (Tekle et al., 2012; Tekle et al., 2016) and in 11 small foci across 5 countries in OEPA (World Health Organization, 2017). These successes led in 2012 to the decision to move from ensuring sustainable systems for control of onchocerciasis as a public health problem to elimination of onchocerciasis from 80% of endemic African countries by 2025; i.e., changing the focus from reducing the disease burden to permanent interruption of transmission in all areas.

This change means that countries face a number of new challenges including, but not limited to, delineating all areas that now need to be included in interventions, including areas of low prevalence, evaluating progress towards elimination using appropriate epidemiological and entomological evaluations, and, crucially, making decisions on when and where to stop MDAi that take into account the need for collaborations between intervention projects or units within an individual country and between national intervention programs in neighbouring countries (Boakye et al., 2018; Bush et al., 2018; Rebollo et al., 2018; Unnasch et al., 2018). In 2016, WHO issued new guidelines which define procedures and entomological criteria for stopping MDAi and commencing post-treatment surveillance, and for subsequent verification of elimination of onchocerciasis and the start of post-elimination surveillance (World Health Organization, 2016). These guidelines also suggested a definition of an onchocerciasis transmission zone as the epidemiological unit within which these procedures and criteria should be applied.

In the context of onchocerciasis, elimination is “the reduction to zero of the incidence of infection caused by a specific agent in a defined geographical area as a result of deliberate efforts” (Dowdle, 1998) with post-elimination measures based on the outcomes of continued surveillance. Assuming these criteria are appropriate and have been met within the geographic area evaluated, there are, in principle, two different reasons elimination may nevertheless not have been achieved or may not be sustainable (Hedtke et al., submitted). First, residual infections not detected during the evaluations may be sufficient for the ‘locally endemic’ parasite population to recover. Second, infected individuals and/or infective vectors may migrate from areas of active transmission into areas where interventions were stopped, and lead to re-establishment of a parasite population via immigration. The magnitude of this risk depends on the alignment of the geographical boundaries for evaluating epidemiological and entomological indicators of transmission with the boundaries within which parasites (in humans or vectors) move and are transmitted, i.e., the parasite transmission zone. The 2016 WHO guidelines define a transmission zone as “a geographical area where transmission of *O. volvulus* occurs by locally breeding vectors and which can be regarded as a natural ecological and epidemiological unit for interventions”. The experience from the OCP in West Africa has shown that epidemiologically significant transmission occurred over vast areas: re-invasion of geographic areas where larviciding had interrupted larval development by vectors from distant neighbouring areas was observed (Boatin et al., 1997; Dadzie et al., 2018) and demonstrated that long-range transmission poses a significant threat to elimination even if it is at low frequency. Thus decisions on the ‘natural ecological and epidemiological unit for interventions’ need to take into consideration the long-term spatial density and migration patterns of both the vector and the human host (Barrett et al., 2008; Blouin et al., 1995; Criscione and Blouin, 2004; Criscione et al., 2005; Jarne and Théron, 2001; Nadler, 1995; Prugnolle et al., 2005). For onchocerciasis elimination programs to make evidence-based decisions on transmission zone boundaries, they need tools to generate this evidence. We have embarked on a research program to develop such tools.

Rapid Epidemiological Mapping of Onchocerciasis (REMO; based on nodule palpation (Noma et al., 2002)), shows that Africa is a complex mosaic of contiguous regions of low, moderate and high infection intensity (hypo- meso- and hyper-endemic, respectively; Figure 1) (African Programme for Onchocerciasis and World Health Organization, 2013). This continuous but heterogeneous pattern of endemicity suggests that parasite transmission occurs in a similarly continuous mosaic of overlapping “zones” of high, moderate and low transmission. These zones can be delineated using parasite population structure as a proxy for transmission zone boundaries: parasites from the same transmission zone are able to interbreed and thus are more closely related genetically than they are to parasites from different transmission zones, with which they are less likely to interbreed and thus are genetically less related. Consequently, the likelihood that parasites originate from the same transmission zone and the likelihood of parasite transmission between two locations (invasion) can be inferred from the degree of genetic relatedness between them, and defined quantitatively by estimating parameters of population structure (Archie et al., 2009; Real and Biek, 2007; Schwabl et al., 2017).

**Figure 1.**
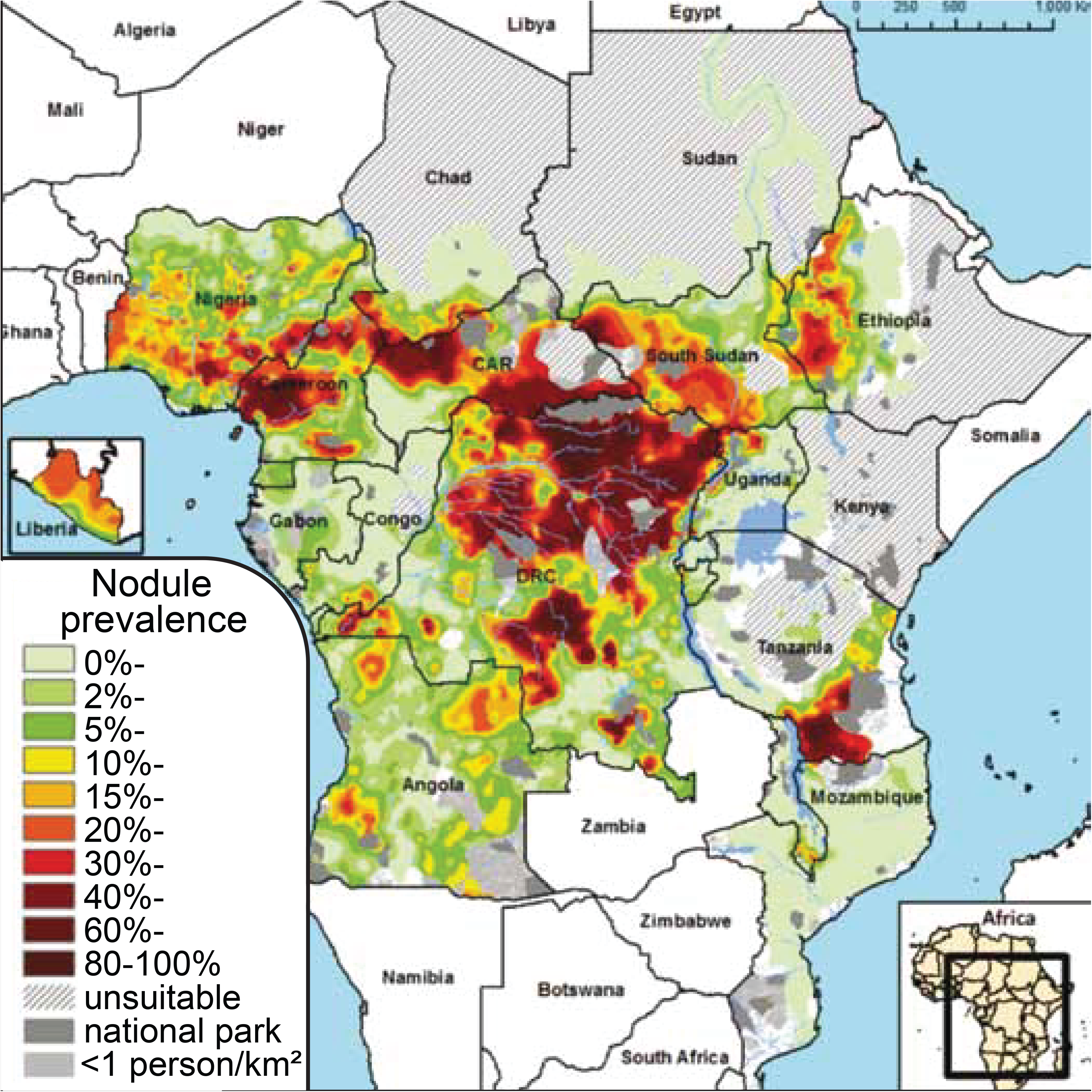
Rapid epidemiological mapping of onchocerciasis (REMO) based on prevalence of nodules across communities with a history of onchocerciasis and covered by APOC. Figure from African Programme for Onchocerciasis Control and World Health Organization (2013).

Early studies of *O. volvulus* found little genetic diversity between parasites isolated from different locations in Africa (Keddie et al., 1998; Zimmerman et al., 1994). However, recent whole genome studies identified large numbers of single nucleotide polymorphisms (SNPs) across the parasite nuclear, mitochondrial, and endosymbiotic bacterial (*Wolbachia* sp.) genomes, with extensive population genetic structure over large (>2000km) (Choi et al., 2016; Crainey et al., 2016) and moderate (>200km) geographic distances (Doyle et al., 2017). However, these genomic studies either focused on only a small number of samples, used pooled samples, or analysed only a small region of the genome. To date, there has been no large-scale survey of genetic diversity and population structure of *O. volvulus* at multiple spatial scales.

In the study reported here, we tested the utility of population genetic markers based on variation in the whole mitochondrial genome of *O. volvulus* for identifying transmission zones and developing genetics-based tools for onchocerciasis control and elimination programs. The generally high copy number of the *O. volvulus* mitochondrial genome makes it easier to amplify than the nuclear genome, and the lack of recombination makes it a useful marker for understanding population demographic history (e.g.,(Avise et al., 1987)). The use of the whole mitochondrial genome, rather than a specific gene, is important for avoiding ascertainment bias and thus underestimating genetic diversity and overestimating gene flow (Ingman et al., 2000; Torroni et al., 2006; Wakeley et al., 2001). We performed high throughput sequencing of the whole mitochondrial genome of 189 adult nematodes sampled from three countries in West Africa (Côte d’Ivoire, Ghana, and Mali) at two geographical scales. Spatial genetic structure between the three West African countries, and within one country, were used to propose genetically-defined transmission zones at the two spatial scales examined.

## 2. Materials and Methods

### 2.1 Sampling

Genomic DNA from 156 individual adult worms was used for long-range PCR, and 36 mt genome sequences were extracted bioinformatically from WGS (S. Hedtke, pers. comm.), for a total of 192 samples. Two of the 192 samples (ASU-7-15 and NLG-86-12) sequenced poorly and were excluded from the analysis, as was one sample (COT-34) that had been sequenced in duplicate. Of the 189 remaining samples, there were two adult male and 187 adult female worms. All adults were obtained by surgical excision of nodules: 13 from 1 sampling location in Côte d’Ivoire, 10 from 4 sampling locations in Mali, and 166 from Ghana (Figure 2). Sampling within Ghana occurred in 16 villages (Table S1, Supplementary Information) across three river basins, with a total of 74 samples from 6 villages in the Black Volta/Tombe (BVT) river basin, 22 samples from 4 villages in the Pru river basin, and 70 samples from 6 locations in the Daka river basin. Individual worms were isolated from nodules using collagenase digestion (Schulz-Key, 1988). Samples from the same river basin or country location are subsequently referred to as 'sample sets'. Details of the collection of the Ghanaian samples have been reported previously (Ardelli and Prichard, 2004, 2007).

**Figure 2.**
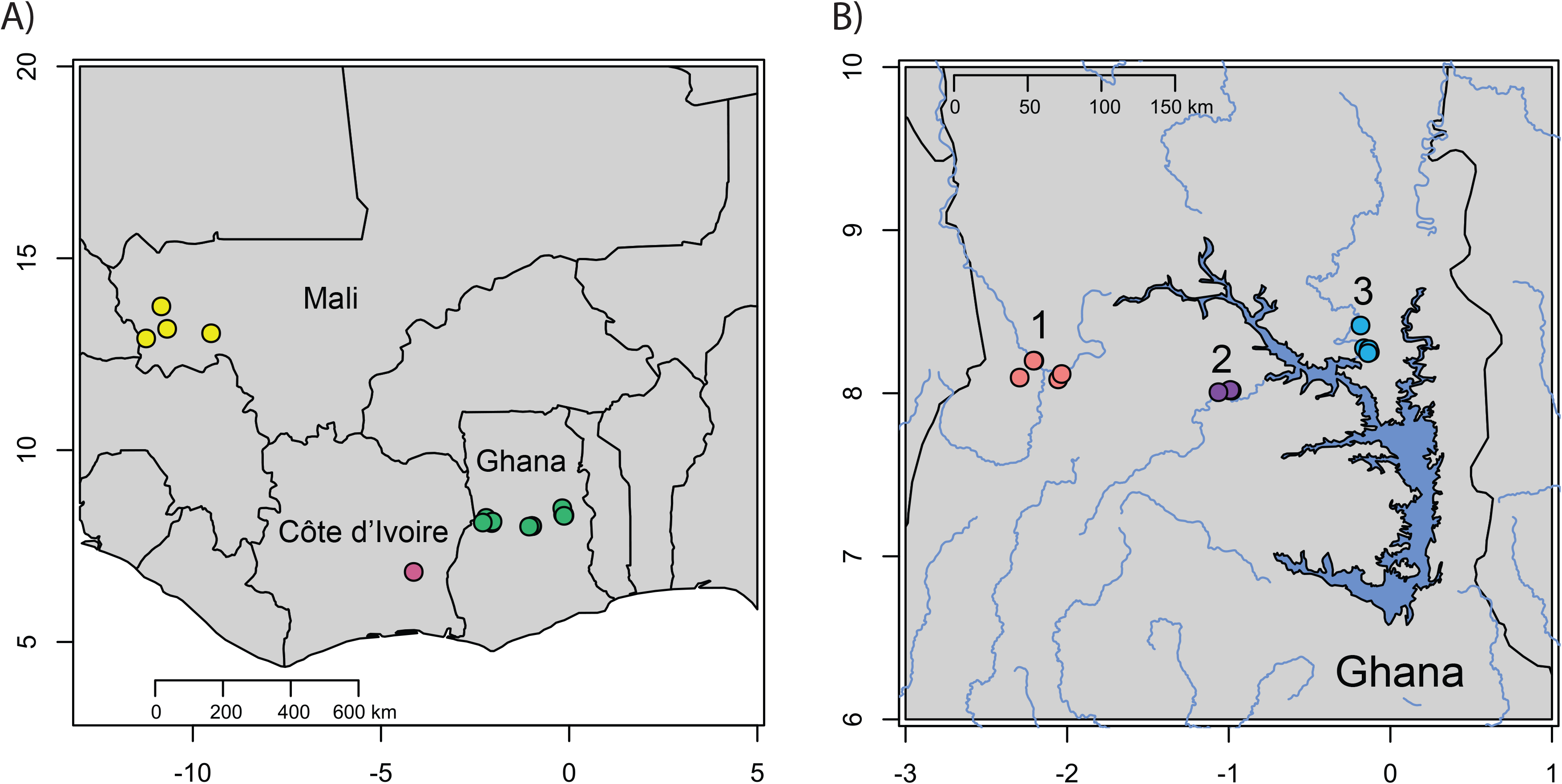
Sampling locations for *O. volvulus* collected. A) Sampling across the African countries of Mali (yellow), Côte d’Ivoire (mauve), and Ghana (green). B) Sampling across the Black Volta/Tombe (red), Pru (purple), and Daka (blue) river basins within Ghana.

### 2.2 DNA extraction and PCR Amplification

DNA was extracted from each worm using the DNeasy^®^ Tissue Kit (Qiagen, Hilden, Germany) following the manufacturer’s instructions. Whole mitochondrial genomes from each nematode sample were PCR amplified using one of three long range PCR strategies: (i) as a single amplicon of 13467 bp, 23 individuals (ii) as two amplicons of 6870 bp and 6912 bp, 95 individuals, or (iii) as three amplicons of 5171 bp, 5572 bp, and 5838 bp, 36 individuals (Figure 3; see Table S2 for primer sequences). PCRs using the GoTaq Long PCR Master Mix (Promega Corp, Wisconsin, USA) using 0.6-0.8 μM of each primer and a modified touchdown PCR protocol for the largest amplicon (Don et al., 1991) (Supplementary Information, Table S3). Each reaction was visualised on a 1 % agarose gel using a 1 kb DNA ladder (New England Biolabs) and a Lambda ladder (Roche) to confirm product size and specificity.

**Figure 3.**
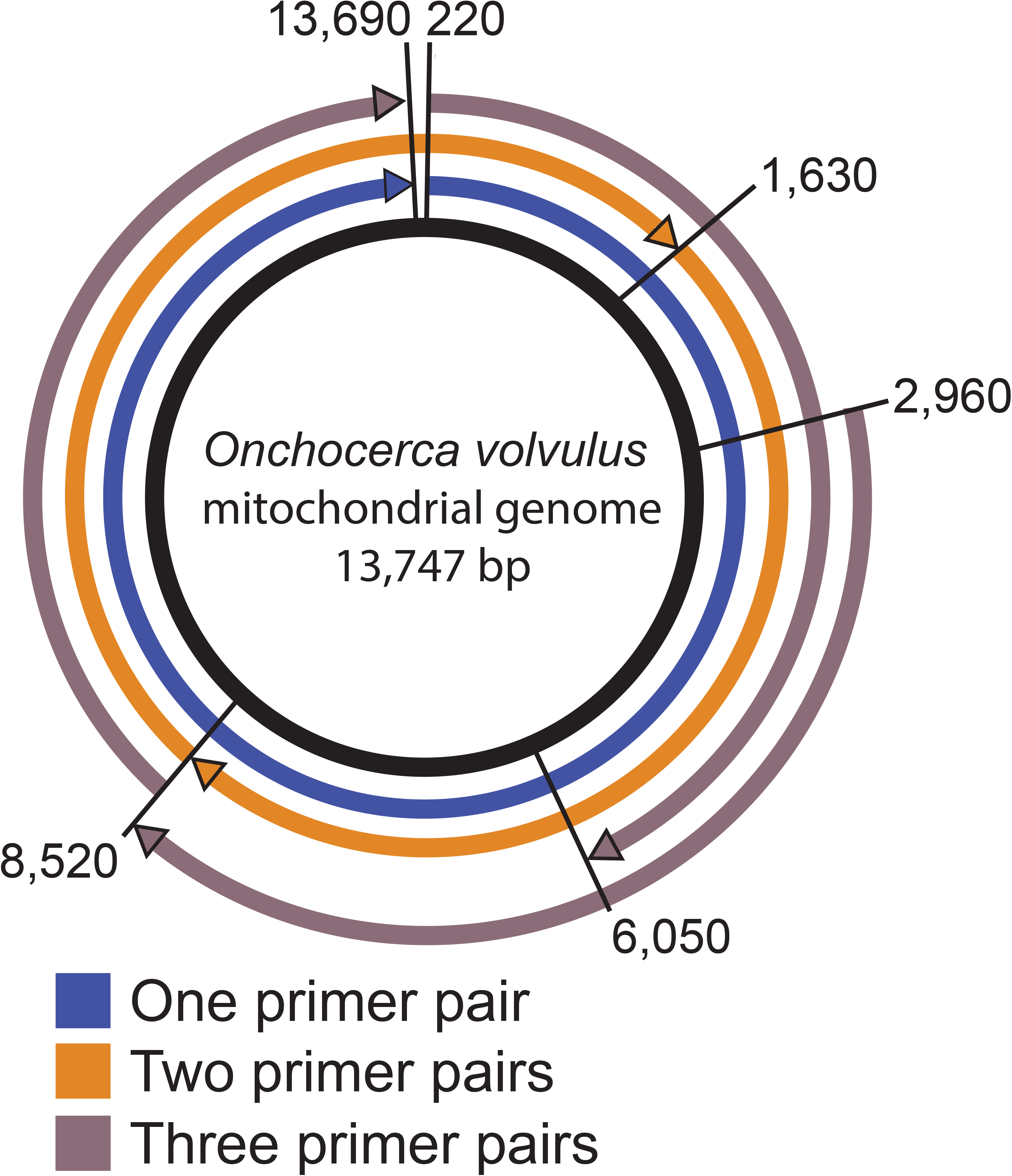
Design for amplification of *Onchocerca volvulus* mitochondrial genome using three possible primer combinations.

### 2.3 Mitochondrial Amplicon Re-sequencing

All PCR products were standardised to a concentration of 500 ng in 50 μl. When an individual sample had multiple PCR products (i.e., the two and three amplicon strategies), the products were combined at equimolar amounts to a total of 500ng in 50ul. Each standardised sample was sheared using a BioRuptor UCD-200 (Diagenode) at high power for 3 minutes with an interval of 30 seconds shearing and 30 seconds rest. Library preparation was performed using a modified Illumina TruSeq DNA HT Sample Prep kit protocol (https://nematodegenetics.files.wordpress.com/2014/05/truseq-ht-workflow-sept2013.docx) to allow Agencourt AMPure XP bead (Beckman Coulter) clean-up in a plate to minimise DNA loss from plate transfers. Samples were then standardised to 40 ng using a Qubit^®^ 2.0 Fluorometer (Invitrogen Life Technologies). Each standardised sample was combined and cleaned using Agencourt AMPure XP beads. The library was size selected for a 400-600 bp insert using Pippin Prep™ (Sage Science) and then sequenced on an Illumina MiSeq^®^ using v2 2×250bp chemistry.

Raw sequence reads had low-quality bases (below quality of 5) and any TruSeq3 Illumina adapters trimmed using Trimmomatic v. 0.32 palindrome trimming (minimum length = 100, 4-bp sliding window with average quality >20, palindrome clip threshold of 30, simple clip threshold of 10). Trimmed reads were then mapped to the *O. volvulus* mitochondrial reference genome (GenBank accession number NC_001861) using BWA-MEM v. 0.7.12 ((Li, 2013); see Supplementary Table S4). The mapping files were sorted and converted to bam files using SAMtools v. 1.2 (Li et al., 2009) and indels were realigned using GATK v.3.3 RealignerTargetCreator and IndelRealigner (DePristo et al., 2011). Duplicate reads were removed using Picard v.1.115 MarkDuplicates (http://broadinstitute.github.io/picard/). Variants were called assuming a haploid genome using two different programs, GATK UnifiedGenotyper and freebayes v.0.9.14-17-g7696787 (with minimum allele count of 5; (Garrison and Marth, 2012)). Variants were filtered using VCFtools v.0.1.12a (Danecek et al., 2011) to exclude primer and missing regions, indels, and non-biallelic sites. Variants with a quality below 30 were removed using bcftools v.1.2 (Li, 2011) (https://samtools.github.io/bcftools/). As freebayes outputs multi-nucleotide polymorphisms, these were split using vcflib v.1 vcfallelicprimitives (https://github.com/ekg/vcflib). Concordant variants between UnifiedGenotyper and freebayes were identified, and discordant and singleton variants were excluded using VCFtools. Variants were annotated using SNPeff v3.6c using an invertebrate mitochondrial codon table (Cingolani et al., 2012) and the *O. volvulus* mitochondrial reference genome (GenBank accession number NC_001861).

### 2.4 Statistical Analysis

Files were converted from fasta to phylip and genepop formats using PGDSpider v.2.0.8.3 (Lischer and Excoffier, 2012). Consensus sequences were generated for each nematode sample using VCFtools vcf-consensus. Minimum spanning haplotype networks were generated using PopART v.1.7 (Leigh and Bryant, 2015). Polymorphic sites and haplotype and nucleotide diversity were calculated using DNAsp v.5.10.1 (Librado and Rozas, 2009). The number of haplotypes, Tajima’s D (Tajima, 1989), Fu’s F (Fu, 1997), pairwise φPT, M values (Slatkin, 1991), and Mantel test comparing genetic and geographic distance (Mantel, 1967) were calculated using Arlequin v.3.5.2.2 (Excoffier and Lischer, 2010). Discriminant analysis of principal components (DAPC), membership probabilities, and individual reassignment scores were generated using adegenet v.2.0.1, with the optimal number of PCs retained determined using the optimascore function (Jombart, 2008; Jombart and Ahmed, 2011; Jombart et al., 2010).

To examine changes in effective population size (N_e_) in the Ghana population over time, we produced a Bayesian skyline plot. Best-fit models of sequence evolution for six possible partitions of the sequence alignment of the entire mitochondrial genomes were determined using the Bayesian Information Criterion in PartitionFinder v1.1.1 (Lanfear et al., 2017), using PhyML for tree estimation (Guindon et al., 2010). Best-fit partitioning schemes placed intergenic regions and genes coding for tRNAs and rRNAs in the same partition, under the HKY+I+G model (Hasegawa et al., 1985), with each codon placed in an additional partition; codon 1 with the HKY+I+G model and codons 2 and 3 with the TrN+I+G model (Tamura and Nei, 1993). These models and partitioning scheme were used to produce a Bayesian skyline plot representing changes in effective population size in BEAST 2 (Bouckaert et al., 2014), with trees linked across partitions, estimating kappa with a starting value of 2.0 and a log normal prior, the proportion of invariant sites with a starting value of 0.2 and a uniform prior, and the gamma shape parameter with a starting value of 1.0 and an exponential prior given 4 rate categories. A strict clock model was used, also linked across partitions, with a clock rate of 9.7 × 10^−8^, based on an estimated substitution rate from *Caenorhabditis elegans* (Denver et al., 2000) and assuming one *O. volvulus* generation per year – a potentially problematic assumption, as filarial nematodes are long-lived and have overlapping generations. The number of intervals within which coalescent events could occur was 5 (i.e., as many as 4 changes in effective population size). Two independent MCMC runs were performed for 150 million generations, storing trees and parameters every 1000 generations, and discarding 10% of each run as burn-in. Stationarity and convergence across runs were assessed using Tracer v1.7.1 (Rambaut et al., 2018), requiring the effective sample size (ESS) to be at least 125 for each run and for the two runs to have strongly overlapping marginal density plots and overlapping mean estimates with corresponding 95% highest posterior density intervals. Stationarity of alpha (the parameter that describes gamma, which in turn describes the shape of rate heterogeneity across sites) and the proportion of invariant sites were difficult to achieve for three of these partitions, with ESS values lower than 100 even after 100 million generations. Examination of the trace files indicated that this was being driven by interactions between these two parameters, as large fluctuations in parameter values were linked. When invariant sites make up a high proportion of the data and when variation at other sites is relatively low, as with these sequences, the algorithm vacillates between incorporating invariant sites into the gamma shape parameter or parameterizing invariant sites separately (Brown et al., 2010; Stamatakis, 2016). These fluctuations were solved by increasing the number of rate categories for the gamma estimation from 4 to 5 and removing the proportion of invariant sites as an independent parameter. Tracer v1.7.1 was used to generate Bayesian skyline plots.

## 3. Results

### 3.1 Genetic Diversity

Variants in the whole mitochondrial genome were identified from 189 samples. A total of 486 high quality SNPs that passed stringent filtering were called, comprised of 297 singleton SNPs (i.e., those only present in one individual) and 189 shared SNPs (i.e., those found in at least 2 samples; Table S5, Supplementary Information). These 189 shared SNPs represented 155 haplotypes (i.e., a group of alleles inherited from a single parent) of which only 19 were identical between 2 or more individuals (Table S5, Supplementary Information). This high haplotype diversity of between 0.96 and 0.99 for each sample set was observed in all communities and countries examined (Table 1). However, the corresponding nucleotide diversity was low, ranging from 0.0006-0.0008 per population, due to the small number of polymorphic sites per individual genome (average per sample 1.05-4.10, Table 1).

**Table 1.** Summary statistics including haplotype and nucleotide diversity, and neutrality tests by country and river basin in Ghana. N: number of samples, S: polymorphic sites, H: number of haplotypes, H_d_: haplotype diversity, π: nucleotide diversity.

| Location | N | S (average/<br>sample) | H | $H_d (\pm SD)$ | $\pi (\pm SD)$ | Tajima's D | Fu's F | Fu &<br>Li's D |
| --- | --- | --- | --- | --- | --- | --- | --- | --- |
| Côte d'Ivoire | 13 | 30 (2.31) | 11 | 0.962<br>(0.050) | 0.0006<br>(0.0001) | -0.605 | -2.604 | -0.081 |
| Mali | 10 | 41 (4.10) | 9 | 0.978<br>(0.054) | 0.0008<br>(0.0001) | -1.327 | -1.506 | -0.994 |
| Ghana | 16<br>6 | 174 (1.05) | 139 | 0.997<br>(0.001) | 0.0006<br>(0.0000) | -2.283* | -24.431* | 1.538 |
| *River basin* |  |  |  |  |  |  |  |  |
| *Black Volta/Tombe* | 74 | 136 (1.84) | 70 | 0.998<br>(0.003) | 0.0007<br>(0.0000) | -2.336** | -24.697** | -2.960 |
| *Daka* | 22 | 56 (2.55) | 20 | 0.991<br>(0.017) | 0.0006<br>(0.0001) | -1.758 | -10.345** | -1.956 |
| *Pru* | 70 | 123 (1.76) | 60 | 0.995<br>(0.004) | 0.0006<br>(0.0000) | -2.281** | -24.750** | -2.584 |

\* $p < 0.01$
\*\* $p < 0.001$

As singletons are not easily distinguishable from PCR error and thus would not be informative, statistical analysis was performed only using SNPs found in more than one individual worm. Tajima’s D, Fu’s F and Fu and Li’s D are statistical tests of neutrality (Fu, 1997; Fu and Li, 1993; Tajima, 1989); i.e., they test whether mutations are passed on to progeny randomly or are subject to selection or other forces that restrict random inheritance by progeny. Non-zero values indicate divergence from neutral variation (when mutation and genetic drift are at equilibrium); a positive value suggests balancing selection (multiple alleles are maintained in a gene pool) or a population bottleneck (only a fraction of a population, and thus only a fraction of the genetic diversity within the population contributed to the next generation), whereas a negative value suggests the occurrence of a selective sweep (when individuals with an allele that provides a survival/fecundity advantage have more offspring that inherit that beneficial allele than those that do not, reducing the overall genetic diversity) or a recent population expansion (when common alleles increase because the number of individuals in the population has expanded faster than new mutations can arise) (Tajima, 1989). Tajima’s D values (Table 1) were negative for the samples from Ghana (p < 0.05), whereas those from Mali and those from Côte d’Ivoire were not significantly different from zero, which may be due to the low number of parasites in each of these sample sets (n = 10 and n = 13, respectively).

A positive or negative Fu’s F, and Fu and Li’s D, indicates a divergence from the expected number of alleles in a population: A positive value suggests balancing selection or a population bottleneck, whereas a negative value suggests an excess of alleles consistent with a population expansion or genetic hitchhiking (when genetic variation is physically linked on a chromosome, selection for an allele at one gene can increase the frequency of the “hitchhiker”, and allele that does not impact survival (Fu, 1997). Fu’s F values (Table 1) are consistent with the Tajima’s D values found, with highly statistically significant (p < 0.01) negative values across the Ghana samples. Both Mali and Côte d’Ivoire sample sets had negative Fu’s F values which were not statistically significant (p > 0.05), which again may be due to the low sample size.

The haplotype network shows a star-like pattern of many closely related haplotypes as indicated (Figure 4 A), consistent with the haplotype diversity statistics (Table 1). There is some clustering of haplotypes at a country level: most Côte d’Ivoire and Mali haplotypes group closely together in the networks (are more closely related to each other than they are to Ghanaian haplotypes), whereas Ghanaian haplotypes are spread throughout the whole haplotype network (Figure 4 B).

**Figure 4.**
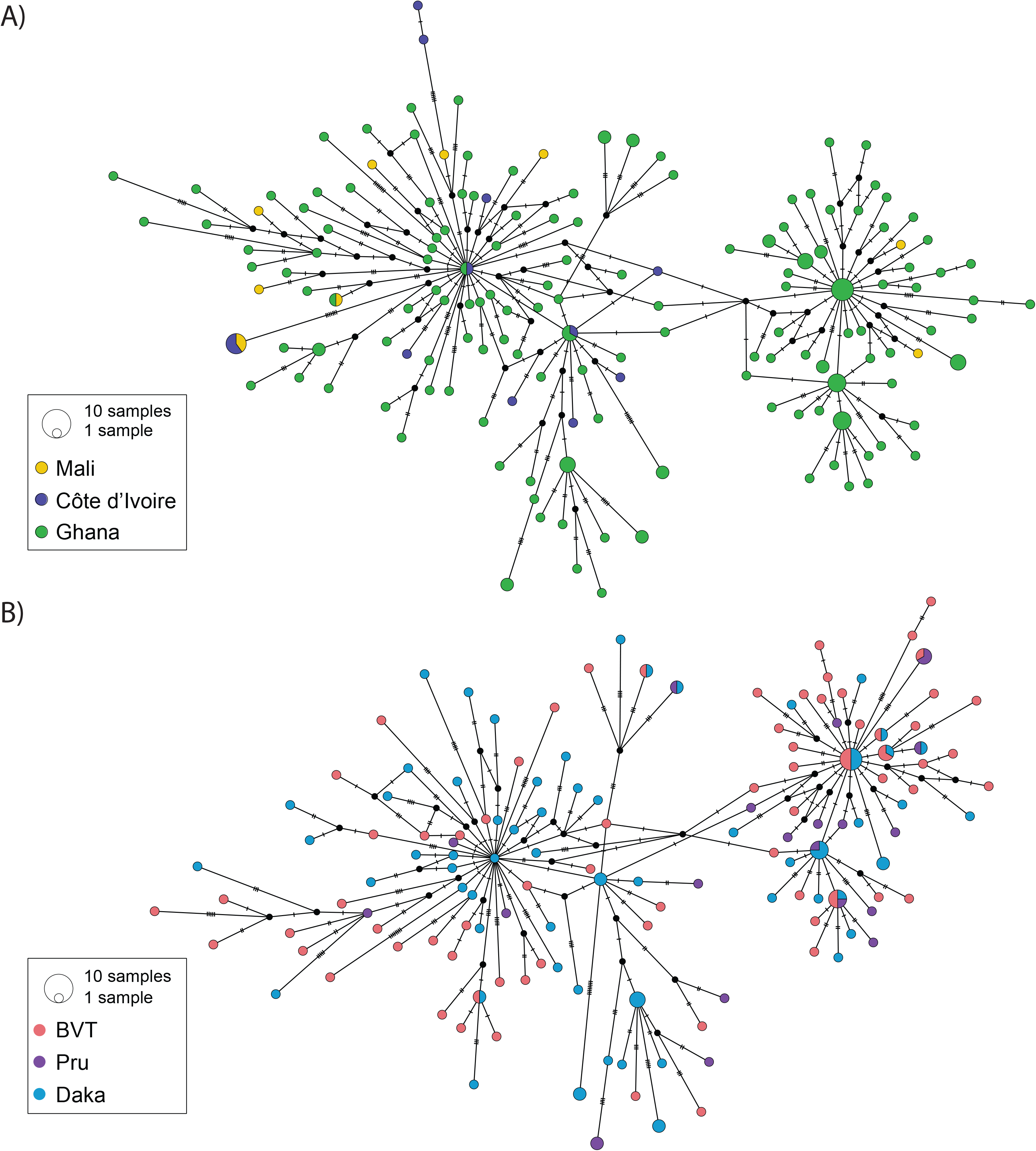
Haplotype networks based on whole mitochondrial genome sequences. A) Haplotype network for *O. volvulus* sampled in three different countries in West Africa: Mali, Côte d’Ivoire, and Ghana; B) Haplotype network for *O. volvulus* sampled across three river basins within Ghana

### 3.2 Population Structure

Population structure was analysed at 3 spatial scales: between countries, between river basins within a single country (Ghana) and between individual villages (Figure 2, Table 2 and Table 3). Pairwise comparisons of genetic sequence variation indicate statistically significant differences between Côte d’Ivoire and Ghana (Table 2; φ_PT_ = 0.0180, p < 0.001) and between Mali and Ghana (φ_PT_ = 0.0112, p < 0.05) but there was no difference found between Côte d’Ivoire and Mali (φ_PT_ = 0, p > 0.05). Consistent with these measures of differentiation, Slatkin’s M values (Slatkin, 1991) indicate a high level of gene flow, i.e., historic or current migration of parasites followed by interbreeding, between Côte d’Ivoire and Mali, and much lower gene flow between Ghana and Côte d’Ivoire, and between Ghana and Mali (Table S6, Supplementary Information). A pairwise comparison of genetic sequence variation by river basin within Ghana (Table 2) showed a low level of genetic differentiation between the parasites from all three river basins (φ_PT_ values were not significantly different from zero; p > 0.05). Corresponding M values for the three river basins (Table S6, Supplementary Information) showed higher gene flow between the Daka and Black Volta/Tombe river basins, relative to the gene flow between the Pru river basin and both the Daka and Black Volta/Tombe river basins. When samples were analysed pairwise by the village within Ghana where they had been collected, no measures of genetic differentiation were significantly different from zero (p > 0.05). Furthermore, M values between most villages suggest a high level of gene flow and thus past or current transmission between all villages (Table S7, Supplementary Information).

**Table 2.**
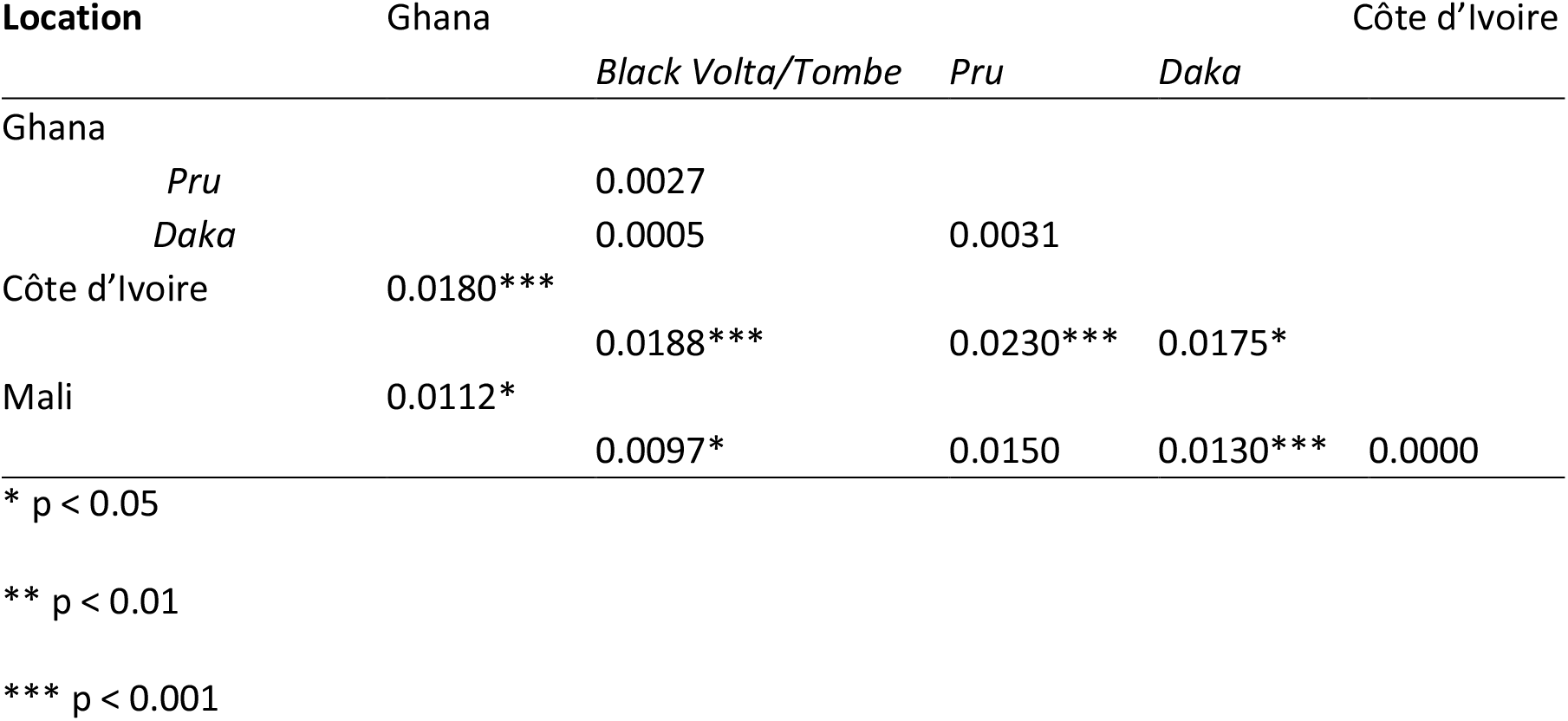
Result of pairwise comparisons of genetic differentiation (φ_PT_) between parasites from Ghana, Côte d’Ivoire and Mali and parasites from three river basins in Ghana (in italics)

**Table 3.**
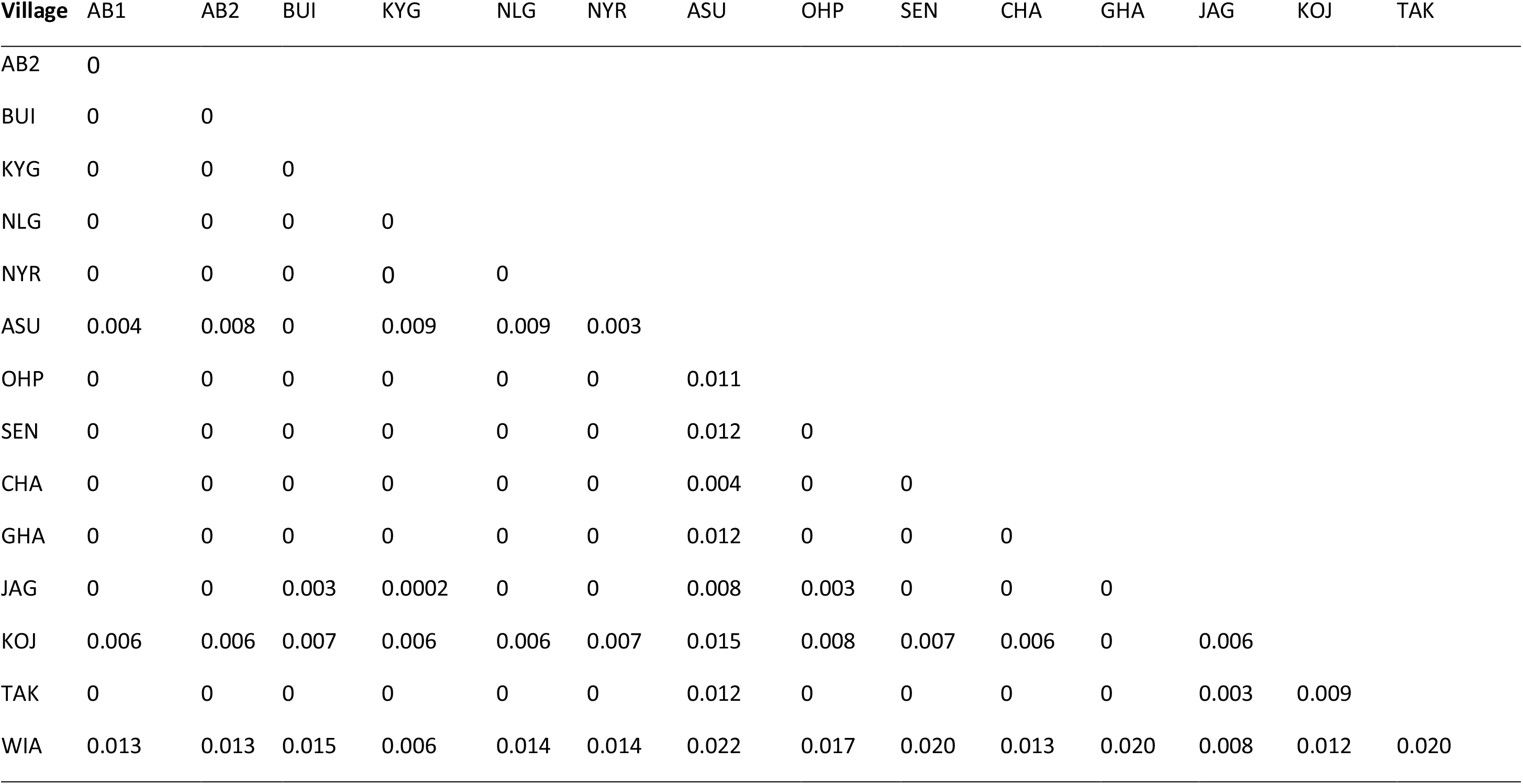
Results of pairwise comparisons of genetic sequence variation (φ_PT_) of between parasites from the inhabitants of 15 villages in Ghana.

| Village | AB1 | AB2 | BUI | KYG | NLG | NYR | ASU | OHP | SEN | CHA | GHA | JAG | KOJ | TAK |
| --- | --- | --- | --- | --- | --- | --- | --- | --- | --- | --- | --- | --- | --- | --- |
| AB2 | 0 |  |  |  |  |  |  |  |  |  |  |  |  |  |
| BUI | 0 | 0 |  |  |  |  |  |  |  |  |  |  |  |  |
| KYG | 0 | 0 | 0 |  |  |  |  |  |  |  |  |  |  |  |
| NLG | 0 | 0 | 0 | 0 |  |  |  |  |  |  |  |  |  |  |
| NYR | 0 | 0 | 0 | 0 | 0 |  |  |  |  |  |  |  |  |  |
| ASU | 0.004 | 0.008 | 0 | 0.009 | 0.009 | 0.003 |  |  |  |  |  |  |  |  |
| OHP | 0 | 0 | 0 | 0 | 0 | 0 | 0.011 |  |  |  |  |  |  |  |
| SEN | 0 | 0 | 0 | 0 | 0 | 0 | 0.012 | 0 |  |  |  |  |  |  |
| CHA | 0 | 0 | 0 | 0 | 0 | 0 | 0.004 | 0 | 0 |  |  |  |  |  |
| GHA | 0 | 0 | 0 | 0 | 0 | 0 | 0.012 | 0 | 0 | 0 |  |  |  |  |
| JAG | 0 | 0 | 0.003 | 0.0002 | 0 | 0 | 0.008 | 0.003 | 0 | 0 | 0 |  |  |  |
| KOJ | 0.006 | 0.006 | 0.007 | 0.006 | 0.006 | 0.007 | 0.015 | 0.008 | 0.007 | 0.006 | 0 | 0.006 |  |  |
| TAK | 0 | 0 | 0 | 0 | 0 | 0 | 0.012 | 0 | 0 | 0 | 0 | 0.003 | 0.009 |  |
| WIA | 0.013 | 0.013 | 0.015 | 0.006 | 0.014 | 0.014 | 0.022 | 0.017 | 0.020 | 0.013 | 0.020 | 0.008 | 0.012 | 0.020 |

A Mantel test comparing genetic diversity and geographic distance indicates that there is a slight positive correlation (r = 0.1989, p < 0.05) between φ_PT_ values and the approximate distance (in km) between the sample sets from the three countries, which indicates isolation-by-distance (IBD) at this large geographic scale. There was no correlation between φ_PT_ values and distance in km between river basins within Ghana (r = 0.0251, p > 0.05), indicating interbreeding of parasites at this smaller spatial scale.

Discriminant analysis of principal components (DAPC (Jombart et al., 2010)), using the 45 principal components (PCs) that contribute most to genetic variation among the samples, demonstrates that variation within the data can be explained by population structuring at a large geographic scale (Figure 5 A) and at a small geographic scale (Figure 6 A) with some overlap between populations. Membership probabilities across the three countries (Figure 5 B-C) calculated by comparing the DAPC predicted country of origin with the actual country of origin show that the DAPC assign 96.4% of parasites from Ghana correctly to country of origin, but only 69.2% of parasites from Côte d'Ivoire and 50% of parasites from Mali assigned correctly. The lower percentages from Côte d’Ivoire and Mali may be due to the low number of parasites sampled and/or the lack of genetic structure between these two countries (as indicated by the parameters described in Table 2). Within Ghana, membership probabilities across the three river basins (Figure 6 B-C) indicate that there is some clustering of parasites by river basin but many parasites were assigned to the incorrect river basin: 25.7% of BVT-originating parasites, 28.6% of Daka-originating parasites, and 45.5% of Pru-originating parasites were not correctly assigned to their river basin of origin based on the genetic variation in their mitochondrial genome.

**Figure 5.**
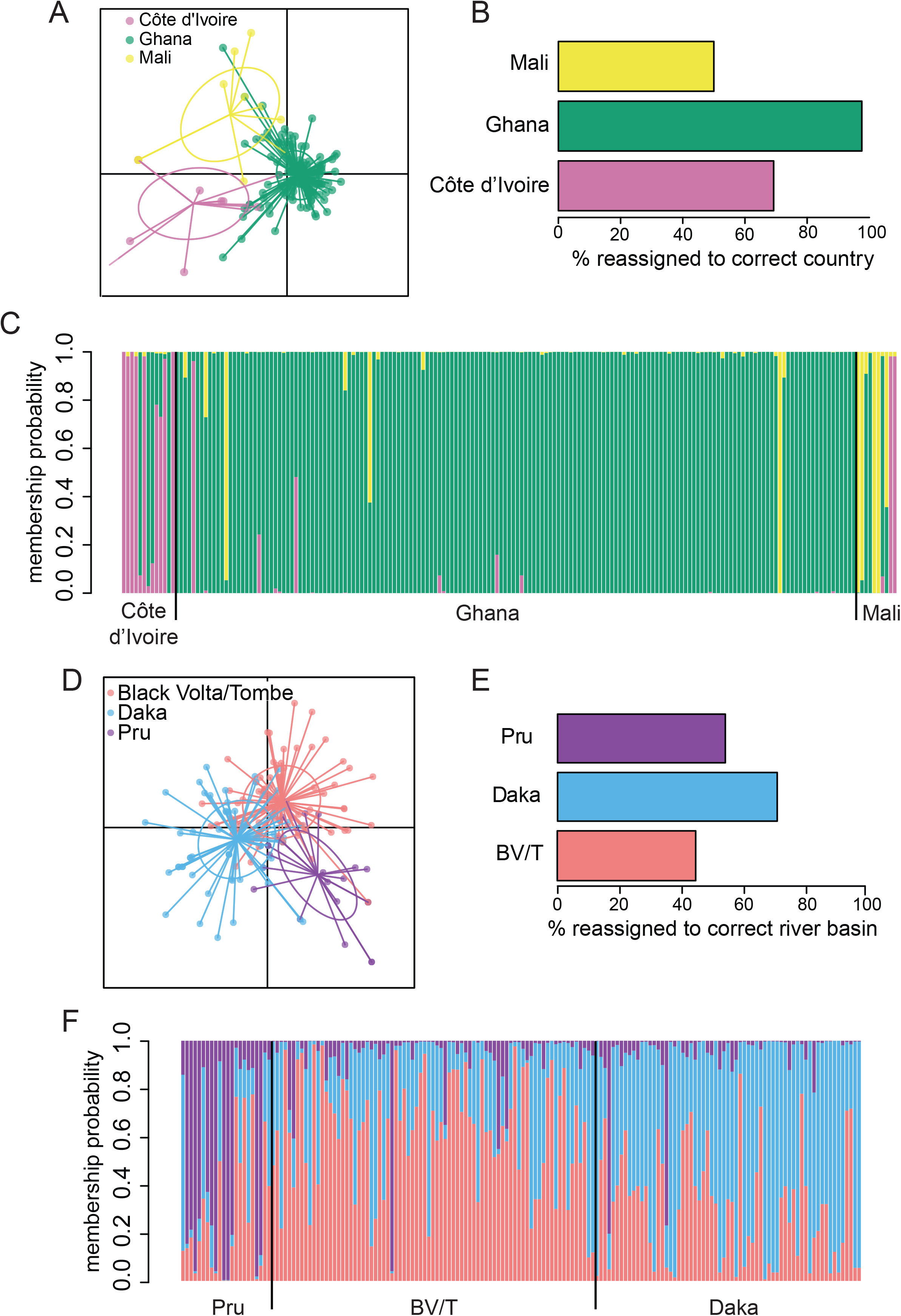
Discriminant analysis of principle components. Top: A) DAPC analysis by country. B) Membership probabilities for parasites from Ghana, Côte d’Ivoire and Mali. C) Percentage of parasites correctly assigned to country of origin. Bottom: D) DAPC analysis by river basins in Ghana. E) Membership probabilities for parasites from river basins in Ghana. F) Percentage of Ghanaian parasites correctly assigned to Ghanaian river basin of origin.

**Figure 6.**
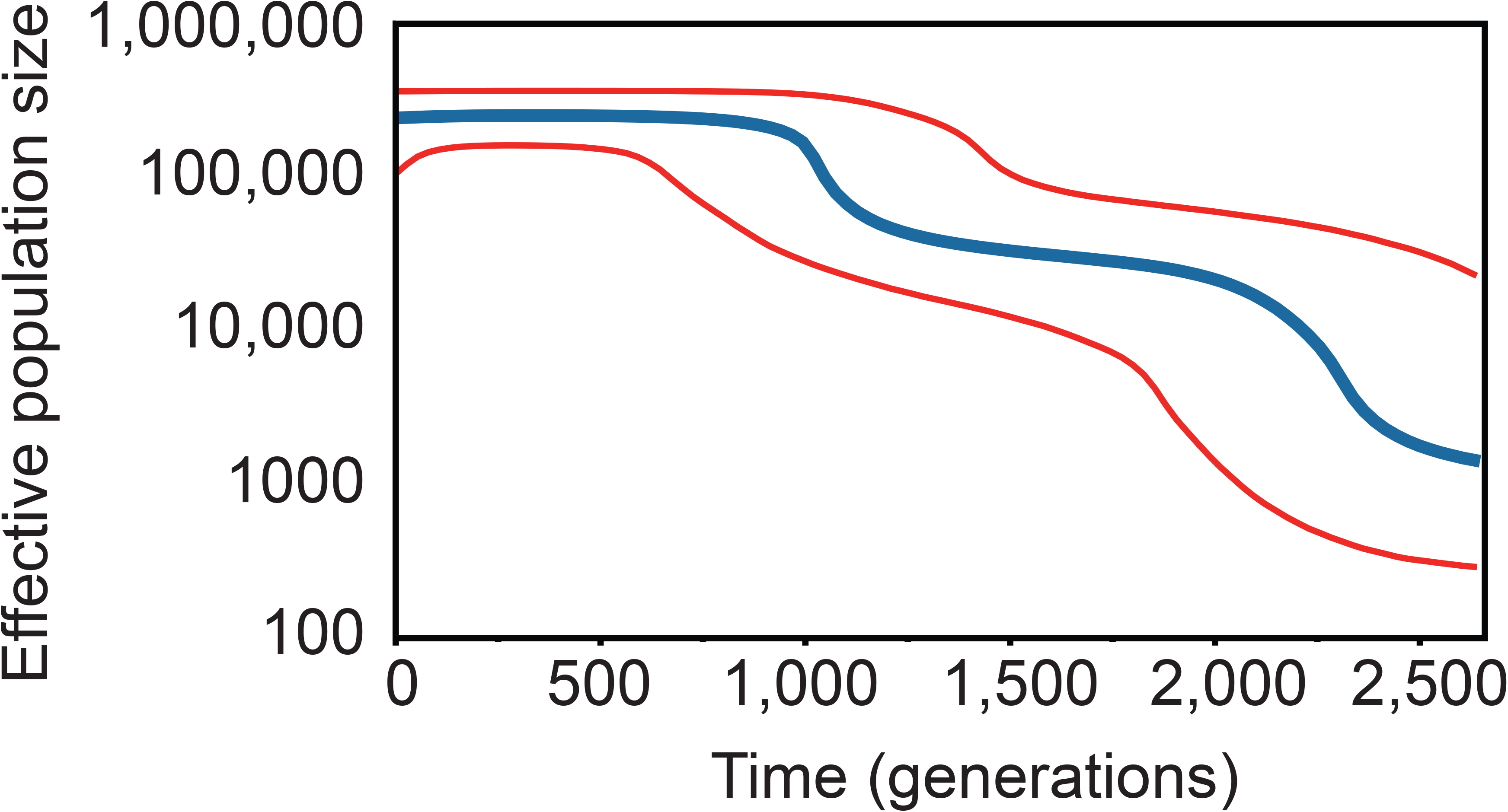
Changes in effective population size over time among West African *Onchocerca volvulus* represented by a Bayesian skyline plot under a coalescent model, assuming one generation per year. Blue line indicates the median estimates, red lines represent the 95% confidence intervals.

## 4. Discussion

### 4.1 Genetic diversity and demographic history of O. volvulus

Reconstruction of the whole genomes of 27 adult *O. volvulus* from Ecuador, Uganda, and forest and savannah regions of West Africa (Choi et al., 2016) had pointed to greater levels of genetic diversity than previously reported (Keddie et al., 1998; Zimmerman et al., 1994). Here, we describe the genetic diversity and population structure of *O. volvulus* derived from complete mitochondrial genomes of 189 adult worms sampled at two spatial scales in West Africa, thus (a) extending significantly the previously available data and (b) analysing population structure at epidemiologically relevant spatial scales for the first time. The 155 unique haplotypes identified in this study (Table 1 & S5, Supplementary Information; Figure 4) demonstrated conclusively that *O. volvulus* populations are extremely genetically diverse, and that the historical population size of this parasite was likely very large. Furthermore, the close relationship between the haplotypes suggests they share an evolutionarily recent origin, consistent with a rapid, large population expansion following an evolutionarily recent host switch (Figure 4). The population expansion most likely occurred after the speciation event that gave rise to *O. volvulus* and *O. ochengi* as a result of proposed host switch by their most recent common ancestor following the introduction of domesticated cattle in Northeast Africa approximately 2,500 – 10,000 years before present (Bain, 1981; Keddie et al., 1998; Lefoulon et al., 2017; Marshall and Hildebrand, 2002). Following the host switch, the new species (*O. volvulus*) presumably expanded rapidly into the naive human population of sub-Saharan Africa, accumulating new mutations in its bottlenecked mitochondrial genome. The large population size (10^6^) suggested by these data is unsurprising when pre-control infection prevalence in the sampled regions is considered (WHO Expert Committee on Onchocerciasis Control (1993 : Geneva and World Health Organization, 1995; World Health Organization and Onchocerciasis Control Programme in West Africa, 1997).

### 4.2 O. volvulus population structure in West Africa

Our data indicate that there was no significant population differentiation between *O. volvulus* from Côte d’Ivoire and Mali (Table 2) which implies that parasites from Mali and Côte d’Ivoire constitute a single population. While investigation of a larger number of parasites from more areas in Mali and Côte d’Ivoire is required, the lack of genetic structure between the parasite populations we examined could be explained as a consequence of the periodic long-range vector migration with the Southwest-Northeast monsoonal winds that required the expansion of the original OCP area (World Health Organization and Onchocerciasis Control Programme in the Volta River Basin Area, 1985; World Health Organization and Onchocerciasis Control Programme in West Africa, 1997). In this context, it is clear that this long-range vector migration is epidemiologically significant and is most likely the primary determinant of the parasite’s population structure and hence of the spatial boundary of a parasite transmission zone, at the geographic scales analysed here.

In contrast, our data indicate that *O. volvulus* populations are genetically structured (i.e., not closely related) between Ghana and Côte d’Ivoire and between Ghana and Mali (Table 2). This implies restricted gene flow east-west between Ghana and Côte d’Ivoire and Southeast-Northwest between Ghana and Mali: parasites in Ghana mate infrequently with parasites from Côte d’Ivoire and Mali. While some *Simulium* spp. subtypes can fly up to 250km (Garms et al., 1982; Post et al., 2013) and up to 500km when aided by favourable wind conditions (Baker et al., 1990; Garms, 1981; Garms et al., 1979; Johnson et al., 1985; Magor and Rosenberg, 1980). The maximum distances separating our sampling locations along the transect between Mali and Ghana, and between Côte d’Ivoire and Ghana are larger (>500-1000km), and do not align with the long-range vector migration between Côte d’Ivoire and Mali that follow the Southwest-Northeast monsoon winds (Baker et al., 1990; Garms and Ochoa, 1979; Walsh et al., 1981; World Health Organization and Onchocerciasis Control Programme in the Volta River Basin Area, 1985). This combination of distance, wind direction, and *Simulium* spp. flight capabilities likely restricts Ghana/Mali and Ghana/Côte d’Ivoire vector movement while facilitating Côte d’Ivoire/Mali migration, and hence also restricts parasite gene flow between Ghana and Cote d’Ivoire and Ghana and Mali, accounting for the observed population structure.

Thus, the parasite population structure we found across the sampling locations in the three countries is consistent with known vector migration patterns and *Simulium* spp. flight capabilities. It is noteworthy that the lack of mitochondrial population structure between Côte d’Ivoire and Mali we found is in contrast to the O-150 population differences between the ecotypes of the sampling locations – forest in Côte d’Ivoire and savanna in Mali (Duke et al., 1966), although we cannot exclude the possibility that diversity levels may have been underestimated due to low sampling size.

We found limited genetic differentiation among parasites from three river basins along as east to west transect of the savannah transition zone in Ghana, suggesting ongoing gene flow and hence parasite transmission, not only within but also between the three river basins sampled. The two furthest river basins (BVT and Daka) are separated by less than the maximum flight range of *Simulium* spp., suggesting that vector movement between river basins is sufficient to maintain parasite transmission and gene flow at this spatial scale. This in turn implies that long-term epidemiological patterns across the three river basins are determined by long-range transmission events rather than short-range focal transmission centred around breeding sites within a given river basin. Movement of infected humans could, of course, also contribute to parasite interbreeding (both within and across countries).

There are some notable exceptions to the lack of population structure between the sampled villages in Ghana. Parasites from inhabitants of Asubende (Pru River basin), Wiae (Daka River Basin) and Kojobone (Daka River Basin) are somewhat differentiated from those obtained from inhabitants of surrounding communities. Although not statistically significant, this weak genetic differentiation may be noteworthy because Asubende is one of the villages where sub-optimal response to ivermectin was first detected phenotypically (Awadzi et al., 2004; Frempong et al., 2016; Osei-Atweneboana et al., 2007). Parasites from Asubende were genetically characterized previously (Doyle et al., 2017), and those with a sub-optimal response phenotype were genetically distinct in nuclear genome comparisons, indicating that the unusual genetic structure of the parasite population from inhabitants of Asubende is not restricted to the mitochondrial genome. We suggest that the unusual genetic profile of Asubende parasites also influences the accuracy of parasite assignment to the Pru basin (which is poor relative to assignment to the Daka and Black Volta basins).

### 4.3 Implications of Population Structure on Delineation of Transmission Zones

A significant advantage of population genetic analysis is that population genetic parameters are an objective and quantitative measure of past transmission between any two locations. Large values for measures of population structure (such as the φ_PT_ statistic for mitochondrial data or the F_ST_ statistic for nuclear genotypic data) imply restricted gene flow (i.e. restricted interbreeding) and hence little or no history of transmission between those locations. Conversely, values for measures of population structure that aren’t statistically significantly different from zero, such as those observed between locations in 3 river basins in central Ghana (Tables 2, 3), imply unrestricted gene flow and hence a history of high transmission between river basins. In other words, population genetic structure is the product of restrictions on gene flow. In parasitological terms, gene flow can only occur when there is parasite transmission between two locations. Thus, the population structure reported here is a measure of the extent of transmission between the geographic areas where the parasites we analysed were collected. Consequently, our data suggest that parasite transmission in this part of West Africa occurs across large distances, consistent with known patterns of vector migration.

The observation that these data are consistent with an isolation-by-distance (IBD) model for parasite population structure and hence parasite transmission is important. Isolation-by-distance population structure implies a continuum of gene flow, and hence implies also a continuum of onchocerciasis transmission, such that locations that are relatively closer together will experience more transmission between them than locations further apart. The important feature of this analysis is the distance and direction over which this transmission occurs. Our analysis indicates that epidemiologically relevant transmission occurs over scales of at least 100 – 200 km, and may occur over greater distances under conducive wind conditions. This latter observation is consistent with the known behaviour of *S. damnosum* in West Africa, but the demonstration of epidemiologically relevant, east-west transmission between river basins in Ghana is novel and has significant implications for the onchocerciasis elimination.

### 4.4 Implications of Population Structure for Decisions to Stop Interventions

A clear implication of our work is that the transmission zones defined by the parasite genetic structure will very likely be much larger than ‘MDAi project areas’ or the focal area surrounding a vector breeding site (the WHO 2016 elimination guideline definition of an epidemiologically relevant transmission zone). For the area covered in this study (central Ghana, Mali and Côte d’Ivoire), transmission zones are measured in hundreds of kilometres and extend over several river basins. It is these large, population genetics-defined transmission zones which should be the units for assessing progress towards elimination and criteria for stopping treatment, and for defining the extent of post-treatment cessation surveillance and estimating the risk of post-treatment cessation recrudescence (Hedtke et al., submitted).

Population genetics-defined transmission zones are based on quantitative measures, which can be used to derive estimates of the risk of post-treatment recrudescence due to immigration from other areas where criteria for stopping treatment have not yet been met. This risk estimate can be considered by the onchocerciasis elimination programs when making decisions to stop treatment in a specific geographic area. If the acceptable risk level is very low, our data would delineate two large transmission zones (North-South between Mali and Côte d’Ivoire, and East-West within Ghana) with little transmission between them. Conversely, cessation of treatment in one location in a single transmission zone while transmission continues in other locations within the same zone would carry a much higher risk, proportional to the extent of transmission between locations. For example, the lack of genetic structure between parasites sampled from three river basins in Ghana, and thus the high probability of transmission between these three river basins, indicates that a very high risk of recrudescence would have to be acceptable for treatment to be stopped in one river basin, even if the prevalence of infective (or infected) vectors meets WHO guideline criteria for stopping treatment, as long as the prevalence in at least one of the other basins does not. Our data illustrate that for elimination to be sustainable, transmission zones cannot be defined by river basin (or by MDAi project area or human infection prevalence patterns), and decisions on stopping treatment may be risky without population genetic data on the extent of interbreeding among river basins/MDAi project areas/REMO endemicity areas, including areas that are geographically very distant.

The current *O. volvulus* transmission models (EpiOncho and OnchoSim) have been used to predict the time to elimination for different levels of pre-intervention endemicity and duration and therapeutic coverage of MDAi (Basáñez et al., 2016; Stolk et al., 2015), and EpiOncho was used to model the time to measurable levels of recrudescence after cessation of MDAi in areas in Senegal and Mali under a range of assumptions concerning residual autochthonous transmission (Walker et al., 2017). Neither model is currently able to model spatial heterogeneity of endemicity or the impact of migration of a defined number of infected/infective vectors or infected people between different areas, i.e., heterochthonous transmission due to long-range movement. We are working to modify EpiOncho to include quantitative population genetic measures and to allow simulation of different transmission zones (McCulloch et al., 2017). Once available and validated, and provided the geographically relevant population genetic data are available, such a model will allow to quantitate the risk associated with stopping interventions in one geographic area meeting WHO guideline criteria when ‘neighbouring’ areas do not.

### 4.5 The utility of population genetics for identifying the origin of parasites detected during post-treatment and post-elimination surveillance

We have shown (Figure 5 A-C, Table 5) that the population genetic data permit identification of the likely geographical origin of a single parasite. In the Comoé river valley in southwest Burkina Faso, where transmission had been interrupted when the OCP ceased larviciding in 1989, *O. volvulus* infection prevalence rates of up to 71% were identified in 2010-2011 and biannual MDAi was implemented. Recrudescence could be attributable in some areas to long-range migration of infected flies from Côte d’Ivoire, although recrudescence due to undetected infections in the Comoé river valley was not excluded (Koala et al., 2017; Koala et al., 2019). With the methods established here, analysis of parasites from Burkina Faso obtained before and after stopping of interventions could establish whether infections/transmission detected now are due to parasites that were missed when decisions to stop larviciding were made, or due to migration of parasites from other endemic regions via vectors or humans. If these analyses indicate a migratory origin, analysis of parasites from potential source areas could furthermore help identify the source and appropriate cost-effective strategies for addressing transmission in the Comoé river valley.

### 4.6 Conclusions

As African onchocerciasis control programs move towards elimination of *O. volvulus* transmission, they need tools for delineating transmission zones throughout which interventions can be stopped, with a defined risk of resurgence they consider acceptable. We have examined the whole mitochondrial genome sequences of 189 adult nematodes to investigate genetic diversity, demographic history and population structure at three spatial scales in West Africa.

The *O. volvulus* mitochondrial genome was found to be more diverse than previously described, with a demographic history consistent with rapid population expansion after a bottleneck likely associated with host switching from cattle to humans. The data support delineation of (at least) two large transmission zones (North-South between Mali and Côte d’Ivoire, and East-West within Ghana). The boundaries between these large transmission zones, and the probability of transmission between any two points within each zone, can be defined by population genetic measures which are the product of historical patterns of transmission and are objective and quantitative. The population genetic parameters required to define a transmission zone can be estimated cheaply and efficiently from genotyping of parasites caught during entomological evaluations with appropriate sampling. The genetic polymorphisms obtained may be used to estimate the risks of resurgence associated with stopping interventions in areas where stopping criteria are met, when they are not met in other geographic areas, and to determine the origin of suspected post-intervention recrudescence. Population genetic measures can therefore be the tools elimination programs need to inform their decisions about cessation of MDAi and/or the frequency and strategy for post-MDAi surveillance.

The data presented here are one of the largest samples of high-resolution population data for any helminth, but are restricted to a relatively small segment of the sub-Saharan range of onchocerciasis. The methods we used can be readily applied to identify *O. volvulus* genetic variation and population structure across Africa to provide additional data to elimination programs facing the decision on where and when stop interventions.

## Acknowledgements

The authors wish to thank Andrew Robinson for his assistance with data analysis using the LIMS High Performance Computing Cluster, and the members of the Grant lab for their assistance and support with laboratory experiments and data analysis.

